# CD72-semaphorin3A axis; a possible new player in immune regulation

**DOI:** 10.1101/2021.08.24.457482

**Authors:** Eiza Nasren, Sabag-D Adi, Kessler Ofra, Jones Yunens, Neufeld Gera, Vadasz Zahava

**Affiliations:** The Proteomic Unit, Bnai Zion Medical Center; Haifa, Israel; Cancer research center, The Rappaport Faculty of Medicine, Technion, Israel Institute of Technology; Haifa, Israel; Division of Structural Biology (STRUBI), Nuffield Department of Clinical Medicine; Oxford, UK

**Author notes:** Corresponding author: Vadasz Zahava, E. mail.

## Abstract

Semaphorin3A (sema3A) inhibits the activity of B and T cells in autoimmune diseases such as Systemic Lupus Erythematosus (SLE). We have now found that CD72 functions as a novel sema3A binding and signal-transducing receptor. These functions of CD72 are independent of the known sema3A receptor neuropilin-1 (NRP-1). We find that sema3A induces the phosphorylation of CD72 on tyrosine residues and the association of CD72 with SHP-1 and SHP-2. In contrast, sema4D/CD100 inhibits these functions. sema3A signals mediated by CD72, inhibit the phosphorylation of STAT-4 and HDAC-1 and induce the phosphorylation of p38-MAPK and PKC-theta in B-cells derived B-lymphoblastoid (BLCL) cells lacking NRP-1 expression, and in primary B-cells isolated from either healthy donors or SLE (Systemic Lupus Erythematosus) patients. We have also generated a modified truncated sema3A (T-sema3A) which cannot signal via NRP-1 yet still activates inhibitory CD72 signaling. We propose that T-sema3A may have potential as a possible therapeutic for autoimmune diseases such as SLE.

**One Sentence Summary:** CD72 found as a novel sema3A receptor transduces inhibitory signals in Bcells. A modified sema3A can be used to treat autoimmunity.

## INTRODUCTION

For more than three decades, many studies indicated that T regulatory cells (Tregs) are the main suppressive regulators of autoimmune diseases and autoimmunity *(1)*. However, it has recently become clear that B regulatory cells (Bregs) also function as important inhibitors of autoimmune responses. Bregs were initially characterized as CD19^+^CD38^hi^CD24^hi^ cells and were shown to be highly effective in down-regulating T helper cells in systemic lupus erythematosus (SLE) and other autoimmune diseases *(2)*. In later studies, Bregs were also characterized as CD19^+^CD5^+^CD1d^hi^ positive cells *(3)*. Regardless, Bregs were found to be IL-10 and TGF-β producing cells *(4)*. In addition to these membrane markers, Bregs also express regulatory co-receptors such as FcγRIIB, CD22, L1RB1 and CD72. When activated, some of these co-receptors recruit the tyrosine phosphatase SHP-1, which interacts with their immune-receptor tyrosine-based inhibitory motifs (ITIMs) to fine-tune the threshold of B cell receptor (BCR) signaling *(5-7)* CD72 is a B cell surface protein of the C-type lectin superfamily and functions as a negative regulator of B cell activation *(8)*. CD72 was observed to associate with the SHP-1 and SHP-2 secondary messengers in B cells *(9, 10)*. CD72 knockout mice have less mature B-2 cells and more B-1 cells in peripheral blood. In these mice, B cells were found to be hyperactive and hyper-proliferative. They also displayed increased intracellular Ca^++^ influx following IgM cross-linking *(11)*. CD72 has been defined as a CD5 ligand on B cell surfaces. CD19^+^CD5^+^CD1d^hi^ Bregs induce the expansion of CD4^+^Foxp3^+^ and CD4^+^CD25^+^ Tregs, and when further activated with IL-21 induced the expansion of IL-10-expressing Bregs. When the activities of CD72 or CD5 were inhibited, both IL-10 expressing Bregs and Tregs were down-regulated. These observations suggest that the interaction of CD5 and CD72 plays a critical role in maintaining regulatory immune mechanisms *(12, 13)*. The regulatory role of CD72 in B cells was further assessed in SLE. The expression of CD72 in B cells of SLE patients was found to be significantly lower as compared to healthy individuals and was inversely correlated with SLE disease severity. Furthermore, reduced CD72 expression was associated with the onset of lupus nephritis and with the presence of anti-dsDNA autoantibodies *(14)*.

The class-4 semaphorin CD100/sema4D was characterized as a ligand of CD72. Following its binding to CD72, sema4D induces the dissociation of CD72 from SHP-1, thereby switching off negative signals produced by CD72 *(15)*. BCR signals are significantly inhibited when CD100 is absent or down-regulated, allowing SHP-1 phosphorylation and resulting in B-cell hypo-responsiveness. It was also shown that when CD100-deficient mice aged, marginal zone B cells accumulate and develop high autoantibody levels and autoimmunity *(16)*. Additionally, CD72 signaling can push Btk-deficient B cells to overcome their poor responsiveness to BCR signaling *(17)*. Thus, CD72 functions as a B-cells receptor that regulates B-cells mediated immune responses. CD72 is mainly expressed in Bregs and likely required to maintain self-tolerance and prevent autoimmunity.

Semaphorins were initially characterized as repulsive axon guidance factors *(18)*. They are divided into several subfamilies, of which the seven class-3 semaphorins represent the only semaphorins secreted as soluble factors. Semaphorin3A (sema3A), a member of the class-3 semaphorin sub-family, is a potent immune-regulator, active at all the immune response stages. Sema3A down-regulates autoimmunity by suppressing the over-activity of both T-cells and B-cells in autoimmune diseases *(14)*. We have previously observed that CD72 expression was significantly increased in B cells following stimulation with sema3A, but at that time the mechanism was unclear *(14)*. We hypothesized that sema3A may possibly function as a CD72 ligand. In the present study, we show that sema3A is a specific CD72 ligand and that it’s binding to CD72 induces responses such as the down-regulation of STAT-4 and the up-regulation of PKC-theta in B-lymphoblastoid cells expressing recombinant CD72 but not neuropilin-1 receptors (NRP-1). These secondary messengers are strongly implicated in the pathogenesis of autoimmune responses, suggesting that sema3A administration may benefit patients suffering from autoimmune disorders by restoration of self-tolerance. Based upon these observations, we have also generated an active truncated sema3A (T-sema3A) that is able to signal via CD72 but which unlike wild type sema3A is not able to induce NRP-1 mediated signal transduction.

## RESULTS

### Semaphorin-3A binds to the CD72 receptor

To determine if sema3A is capable of binding CD72, we performed a Co-IP experiment using U87MG-ΔNRP-1-CD72^+^ cells, in which the NRP-1 gene was knocked-out using CRISPR/Cas9 *(19)* and in which recombinant CD72 stably expressed. The expression of NRP-1 and CD72 in these cells and in control U87MG-ΔNRP-1 cells was compared at the mRNA and protein levels with parental U87MG cells, which expre1ss only the NRP-1 receptor (Fig. 1 A&B). These experiments reveal that antibodies targeting the V5 epitope tag of CD72 immunoprecipitated sema3A from U87MG-ΔNRP-1-CD72^+^ but not from U87MG-ΔNRP-1 cells. Control antibodies directed against HA failed to immunoprecipitate sema3A or CD72, indicating that the immunoprecipitation is specific. Taken together, these experiments suggest that sema3A binds specifically to the CD72 receptor (Fig. 1C).

**Figure 1:**
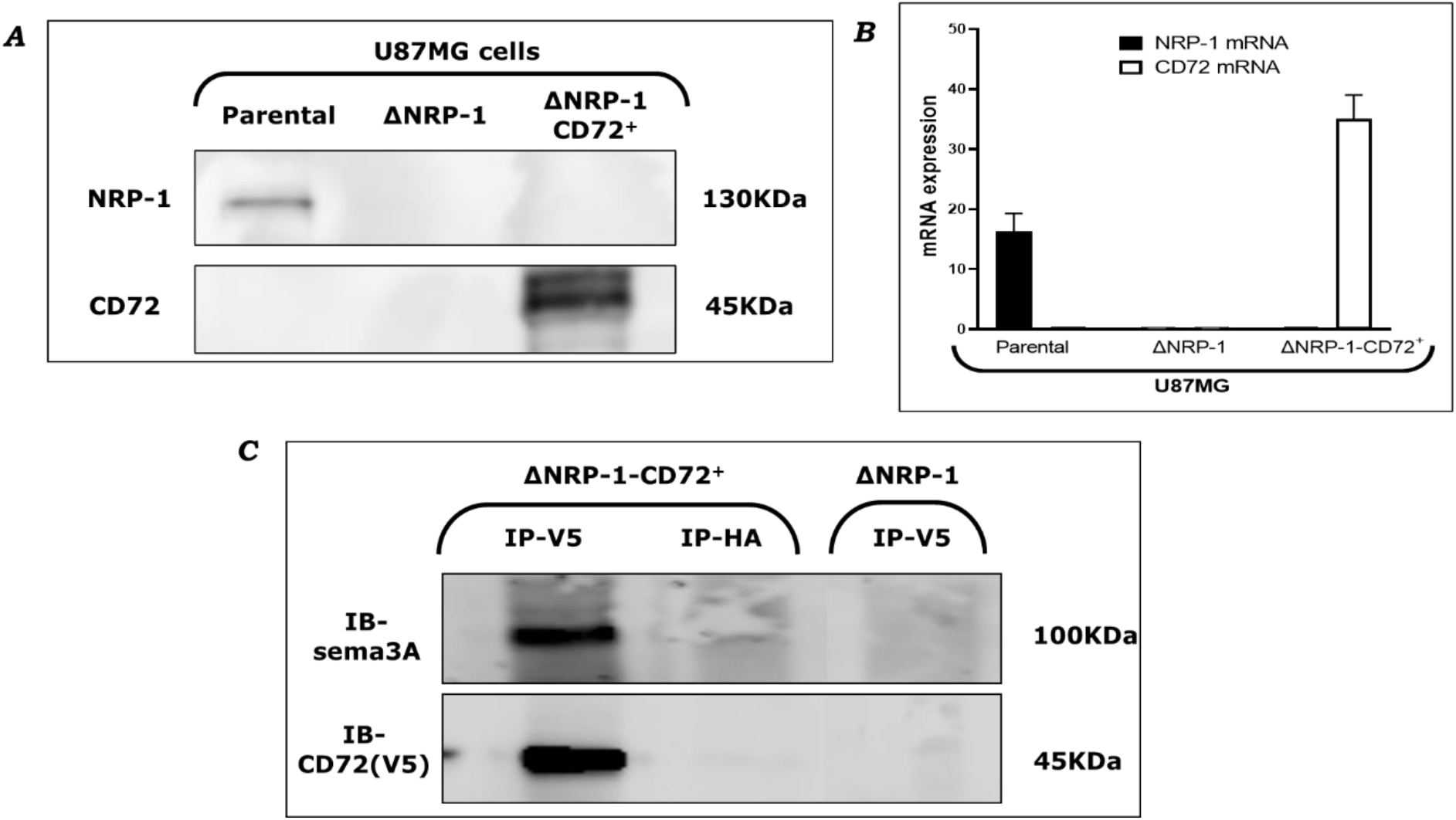
CD72 immunoprecipitates sema3A in U87MG-ΔNRP-1-CD72^+^/V5 cells. **(A)** Western blot showing of CD72 and NRP-1 expression in U87MG, U87MG-ΔNRP-1 and in U87MG-ΔNRP-1-CD72^+^/V5 cell lines. **(B)** Real time PCR of NRP-1 and CD72 mRNA expression in U87MG, U87MG-ΔNRP-1 and in U87MG-ΔNRP-1-CD72^+^/V5 cell lines. **(C)** U87MG-ΔNRP-1 and U87MG-ΔNRP-1-CD72^+^/V5 cells were stimulated with sema3A for 20 minutes at 4°C, followed by 5 minutes incubation at 37°C. Cell lysates were then prepared and immunoprecipitated using antibodies targeting V5 epitope tag (attached to CD72) or HA tag (as a control). Blots prepared from the CD72 immunoprecipitates were probed with antibodies directed against sema3A and CD72 as indicated. Shown are the resulting western blots.

To further verify these observations, we performed a binding assay using sema3A fused to alkaline phosphatase (sema3A-AP). Cells were incubated with conditioned medium containing 5µg/ml of sema3A-AP. Bound sema3A-AP was visualized using BICP/NBT. Sema3A-AP bound to parental U87MG cells expressing NRP-1 as well as to U87MG-ΔNRP-1-CD72^+^ cells, which express CD72 but not NRP-1. In contrast, sema3A-AP failed to bind to U87MG-ΔNRP-1 cells, which lack both receptors (Fig. 2A). These experiments also strongly suggest that CD72 functions as a sema3A binding receptor, and furthermore indicate that sema3A binds to CD72 independently of NRP-1.

**Figure 2:**
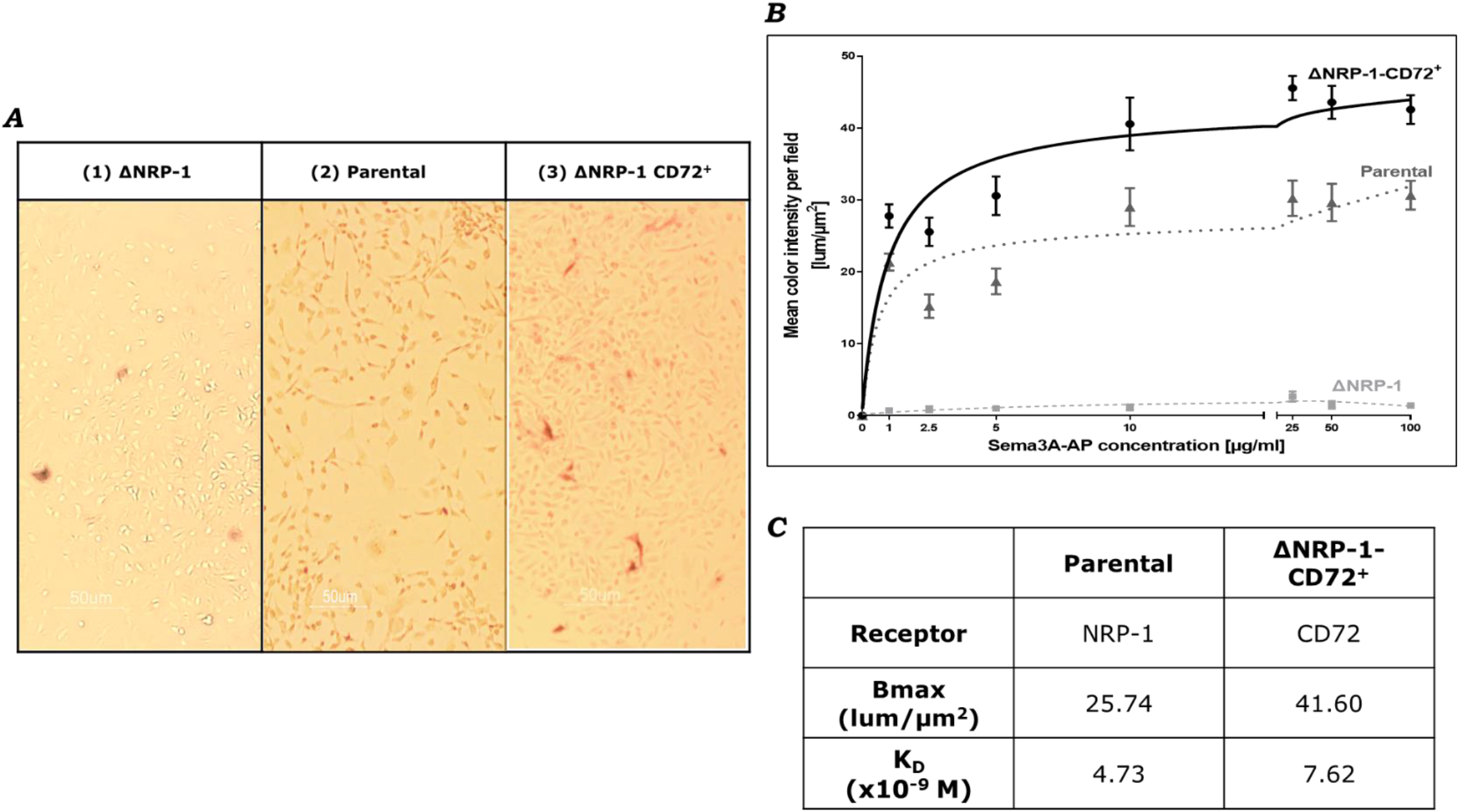
Sema3A –AP binds CD72 receptor. **(A)** Representative images of sema3A-Alkaline phosphatase (AP) binding assay in different U87MG cell lines. Cells were incubated with conditioned medium containing 5µg/ml of sema3A-AP. Bound sema3A-AP was visualized using BCIP/NBT. (1) shows negative staining in U87MG-ΔNRP-1, (2) shows positive staining in U87MG-Parental (Par) while (3) demonstrates positive staining in U87MG-ΔNRP-1-CD72^+^. **(B)** Log-dose binding experiments in which increasing concentrations of sema3A-AP were bound to U87MG cells as well as to U87MG-ΔNRP-1 and U87MG-ΔNRP-1-CD72^+^ cells. Each dot represents the average of the mean color intensity per field (lum/µm^2^) as a function of sema3A-AP concentration (µg/ml) of 6 independent experiments. The graphs represent the best fit binding model for each receptor. (N=6). **(C)** Sema3A binding parameters to NRP-1 receptor in U87MG-par cells and to CD72 receptor in ΔNRP-1-CD72^+^ cells. The one-side total binding equations were calculated using Graph Pad Prism software. KD is the equilibrium dissociation constant. Bmax is the maximal binding in a cell.

In order to determine the dissociation constant for the binding of sema3A to CD72, we performed log-dose binding experiments in which increasing concentrations of sema3A-AP were bound to U87MG cells as well as to U87MG-ΔNRP-1 and U87MG-ΔNRP-1-CD72^+^ cells (Fig. 2B). Sema3A-AP bound to NRP-1 with a somewhat higher affinity than to CD72 (K_D_ ∼4.73×10^−9^ M to NRP-1 versus ∼7.62×10^−9^ M to CD72) (Fig. 2C).

To further examine the kinetics of the sema3A binding to CD72, we performed the binding assay in the presence of competitive inhibitors. Sema3A-AP (5µg/ml) was bound to U87MG-ΔNRP-1-CD72^+^ cells in the presence of increasing concentrations of unlabeled sema3A. Indeed, unlabeled sema3A inhibited the binding of sema3A-AP, displaying a half-maximal inhibitory concentration (IC50) of 9.755µg/ml. Interestingly, a very similar concentration (10.48µg/ml) of sema3A was required for half-maximal inhibition of the binding of sema3A-AP to parental U87MG cells, which express NRP-1 but no CD72, suggesting again that the dissociation constants of sema3A from both receptors are very similar (Fig. 3A). CD72 also functions as a receptor for CD100/sema4D, a semaphorin that does not bind to NRP-1. Indeed, we observed that sema4D was also able to inhibit the binding of sema3A-AP to CD72, suggesting sema3A and sema4D share the same CD72 binding domain (Fig. 3B).

**Figure 3:**
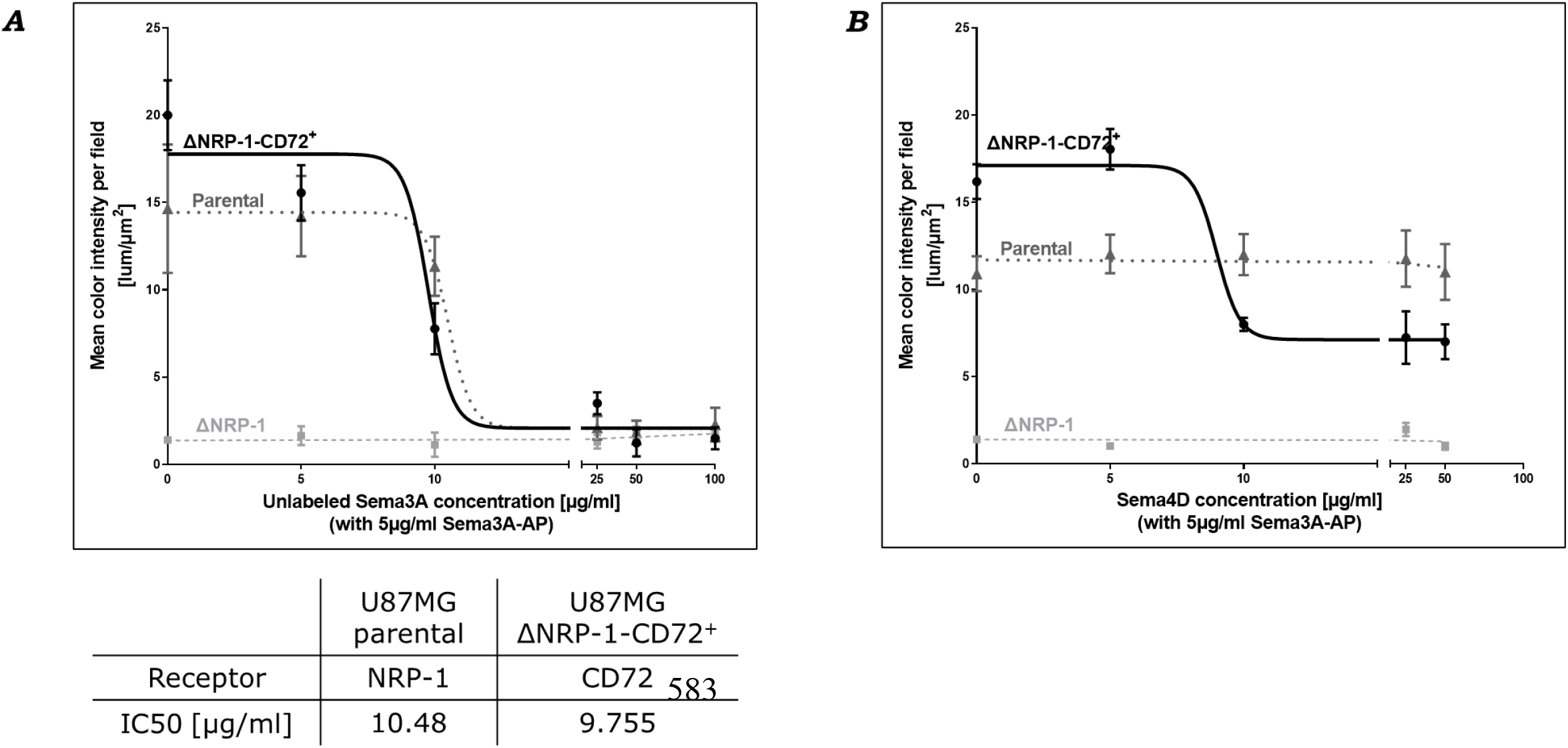
Sema3A and sema4D act as competitive ligands for CD72 receptor. **(A)** Log-dose competitive-binding experiments in which increasing concentrations of unlabeled sema3A with a constant concentration of 5µg/ml of sema3A-AP were bound to U87MG cells as well as to U87MG-ΔNRP-1 and U87MG-ΔNRP-1-CD72^+^ cells. Each dot represents the average of the mean color intensity per field (lum/µm^2^) as a function of unlabeled sema3A concentration (µg/ml) of 4 independent experiments. The graphs shown a similar pattern of inhibition with IC50 of 10.48µg/ml vs. 9.755µg/ml of unlabeled sema3A to NRP-1 and CD72 receptors, in U87MG-Par vs. U87MG-ΔNRP-1-CD72^+^ cells, respectively. (N=4). **(B)** Log-dose competitive-binding experiments in which increasing concentrations of sema4D with 5µg/ml of sema3A-AP were bound to U87MG cells as well as to U87MG-ΔNRP-1 and U87MG-ΔNRP-1-CD72^+^ cells. Each dot represents the average of the mean color intensity per field (lum/µm^2^) as a function of sema4D concentration (µg/ml) of 4 independent experiments. The graphs shown as expected no effect on sema3A binding to NRP-1 and a reduction in binding to CD72. (N=4)

### CD72 transduces Sema3A signals in B cells

In order to find out if CD72 functions as a sema3A signal-transducing receptor in B cells, we used primary B-lymphoblastoid cells (BLCL), which do not express (or almost do not express) NRP-1 or CD72. We over-expressed in these cells the cDNA encoding CD72 to generate BLCL cells expressing CD72 (BLCL-CD72^+^) (Figs. 4A & 4B).

**Figure 4:**
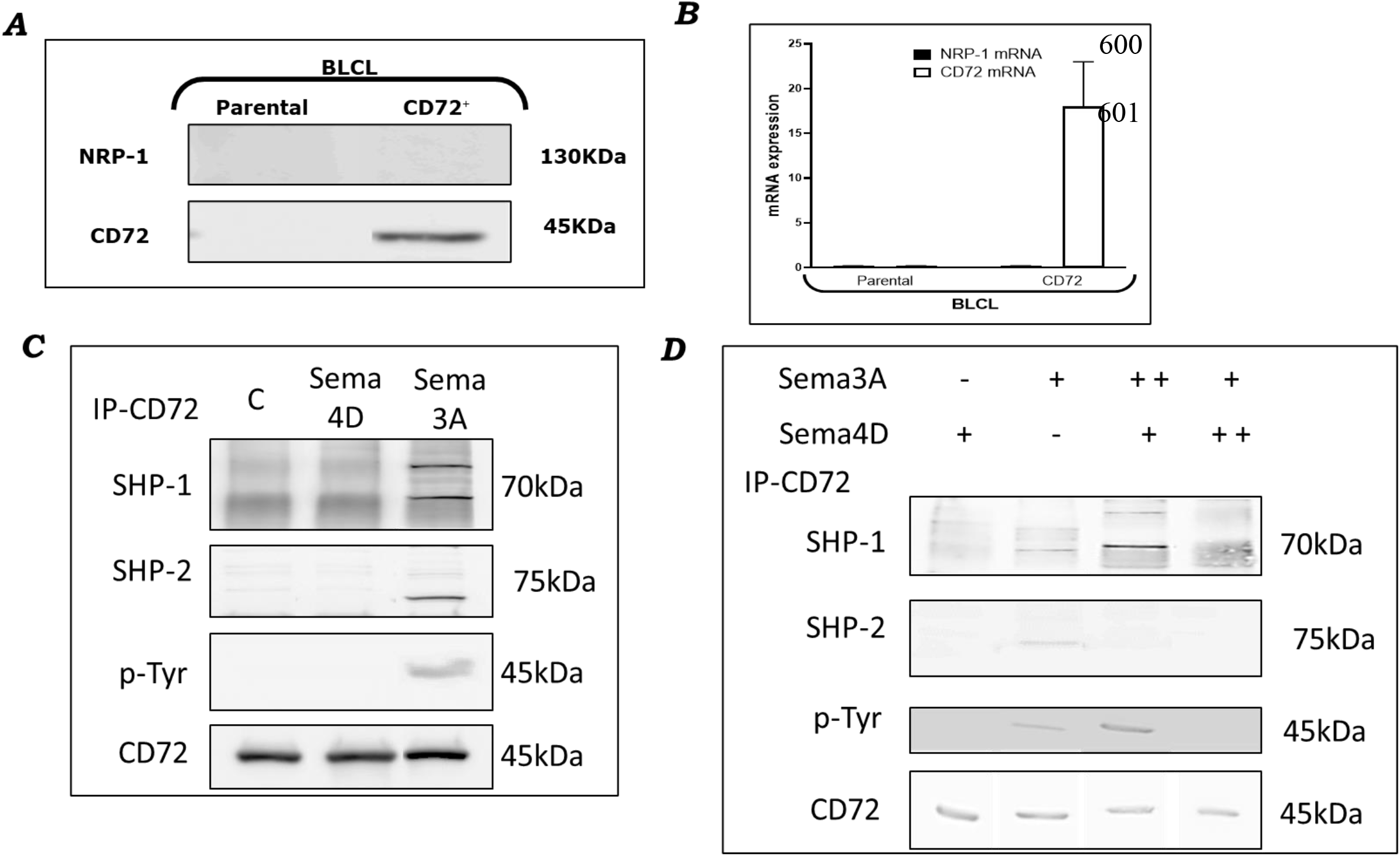
Sema3A promotes the phosphorylation of CD72 and its association with SHP-1 and SHP-2 while sema4D competes with sema3A and inhibits the phosphorylation. **(A)** Western blot showing of CD72 and NRP-1 expression in BLCL and BLCL-CD72^+^ cells. **(B)** Real time PCR of NRP-1 and CD72 mRNA expression in the BLCL and BLCL-CD72^+^ cell lines. **(C)** BLCL-CD72^+^ cells were stimulated with carrier control (C) and either 5ug/ml sema4D or 5ug/ml sema3A for 5 minutes at 4°C. Cell lysates were then prepared and were imunoprecipitated using antibodies targeting CD72. Blots prepared from the lysates or from the CD72 immunoprecipitates were then probed with antibodies directed against SHP-1, SHP-2, phospho-tyrosine or CD72 as indicated. (N=4). **(D)** BLCL-CD72^+^ cells were stimulated with either sema3A or sema4D (+ = 5ug/ml, ++ = 20 ug/ml, respectively) or with a mixture of both semaphorins at the indicated concentrations. Cell lysates were then prepared and were imunoprecipitated using antibodies targeting CD72, and probed with antibodies as in described under C. (N=2). *IC50 is the concentration of agonist that gives a half maximal inhibitory concentration.

CD72 was observed to associate with the SHP-1 and SHP-2 secondary messengers *(9, 10)*. We therefore determined if sema3A is able to induce the association of CD72 with these secondary messengers in BLCL-CD72^+^ cells. Indeed, we found that in contrast to sema4D which was not able to induce the phosphorylation of CD72 and its association with SHP-1 or SHP-2, sema3A induced the phosphorylation of CD72 on tyrosine residues and the association of CD72 with both SHP-1 and SHP-2 (Fig. 4C). Furthermore, because sema4D seems to compete with sema3A for binding to CD72 (Fig. 3B) we determined id sema4D can inhibit the sema3A induced phosphorylation of CD72 and the sema3A induced association of CD72 with SHP-1. Indeed, an excess of sema4D inhibited sema3A induced phosphorylation of CD72 as well as the sema3A induced association with SHP-1 and SHP-2 (Fig. 4D).

We then determined if stimulation of BLCL-CD72^+^ cells with sema3A activates CD72 mediated signal transduction. To identify secondary messengers activated in these cells by sema3A, we used a Full-Moon phospho-explorer-antibody array. Stimulation of BLCL-CD72^+^ cells with sema3A inhibited the phosphorylation of STAT-4 (Tyr693), HDAC-1 (Ser421), Smad-1 (Ser187), and Smad-3 (Ser204) and increased the phosphorylation of p38-MAPK (Tyr182), PKC-theta (Thr538), and FOXO3A (Ser253) (Fig. 5A).

**Figure 5:**
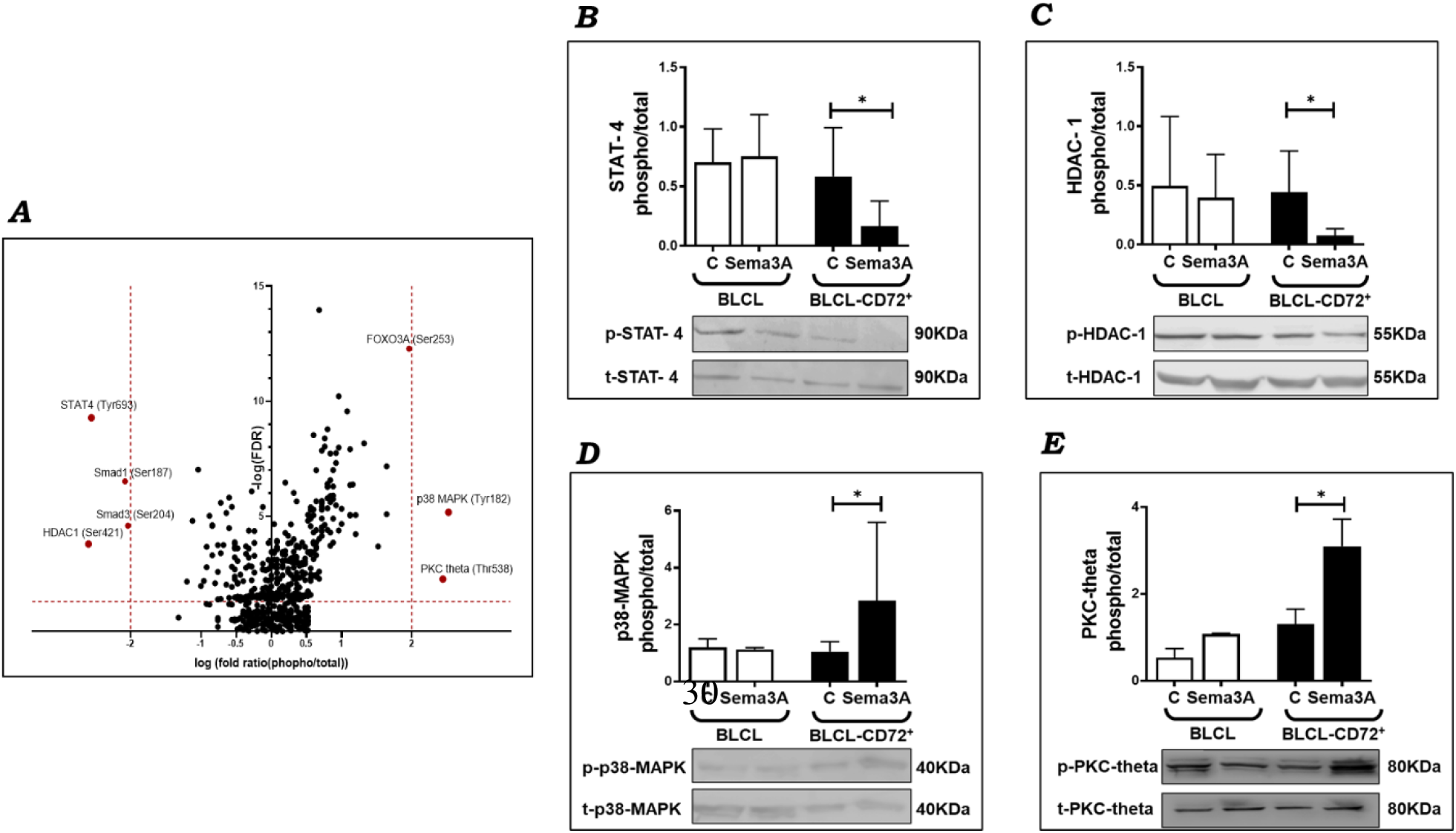
Sema3A binding to CD72 promotes the phosphorylation of p38-MAPK (Tyr182) and PKC-theta (Thr538) and inhibits the phosphorylation of STAT-4 (Tyr693) and HDAC-1 (Ser421) **(A)** BLCL-CD72^+^ were stimulated with 10µg/ml of sema3A for 5 minutes at 37°C (or with carrier control). Then phospho-array experiment was performed on the cell lysates following the Full-Moon antibody array user’s guide, the volcano plot showing the array results. Each dot in the volcano plot represents the log fold change of the phosphorylation statue of signal transduction component (as a result of sema3A simulation compared to control). The components with the most significant change are marked in the plot. **(B)** BLCL and BLCL-CD72^+^ cells were stimulated with 10µg/ml of sema3A for 10 minutes at 37°C (or with carrier control). Cell lysates were prepared in the presence of phosphorylation buffer. Blots prepared from the lysates were then probed with antibodies directed against Phospho-STAT-4 (Tyr693), **(C)** Phospho-HDAC-1 (Ser421), **(D)** Phospho-p38-MAPK (Tyr182) and **(E)** Phospho-PKC-theta (Thr538) as indicated. Shown are representative western blots and the quantification graphs of the mean± SD of the phospho/total ratio of each indicated component from 5 independent experiments.

To verify these results, we performed a phosphorylation assay in which we stimulated BLCL and BLCL-CD72^+^ cells with sema3A followed by western blot analysis using antibodies directed against phosphorylated STAT-4 (Tyr693), HDAC-1 (Ser421), p38-MAPK (Tyr182), and PKC-theta (Thr538). The results reveal no change in the phosphorylation status of pSTAT-4, pHDAC-1, pp38-MAPK, and pPKC-theta in wild type BLCL in response to sema3A. In contrast, as predicted by the phospho-array results, the phosphorylation levels of STAT-4 and HDAC-1 were significantly reduced following stimulation of BLCL-CD72^+^ cells with sema3A (Fig. 5B & 5C), while the phosphorylation levels of p38-MAPK and pPKC-theta were enhanced as predicted following stimulation with sema3A (Fig. 5D & 5E). Taken together, these results suggest that CD72 functions as a high-affinity signal-transducing receptor for sema3A in B cells.

### CD72 signaling modified by sema3A in B-cells of SLE patients

Sema3A changes the phosphorylation state of several secondary messengers following its binding to CD72 in BLCL cell line (Fig.5). To find out if sema3A is able to induce similar changes in B-cells from SLE patients, we purified B-cells from healthy donors and SLE patients. We next compared the expression levels of CD72 and NRP-1 in B-cell from healthy and SLE patients. As previously observed *(14)*, the expression level of CD72 was significantly lower in B-cells of SLE patients as compared with healthy controls, while the expression level of NRP-1 was not significantly different (Fig. 6A). Stimulation with sema3A significantly inhibited the phosphorylation of HDAC-1 (Fig. 6B) and enhanced the phosphorylation of p38-MAPK (Fig.6C). These phosphorylation state changes resemble the changes that were seen in BLCL cells expressing recombinant CD72 (Fig.5). Therefore, it is likely that these changes in the phosphorylation state of these proteins were due to CD72 mediated signal transduction initiated by sema3A.

**Figure 6:**
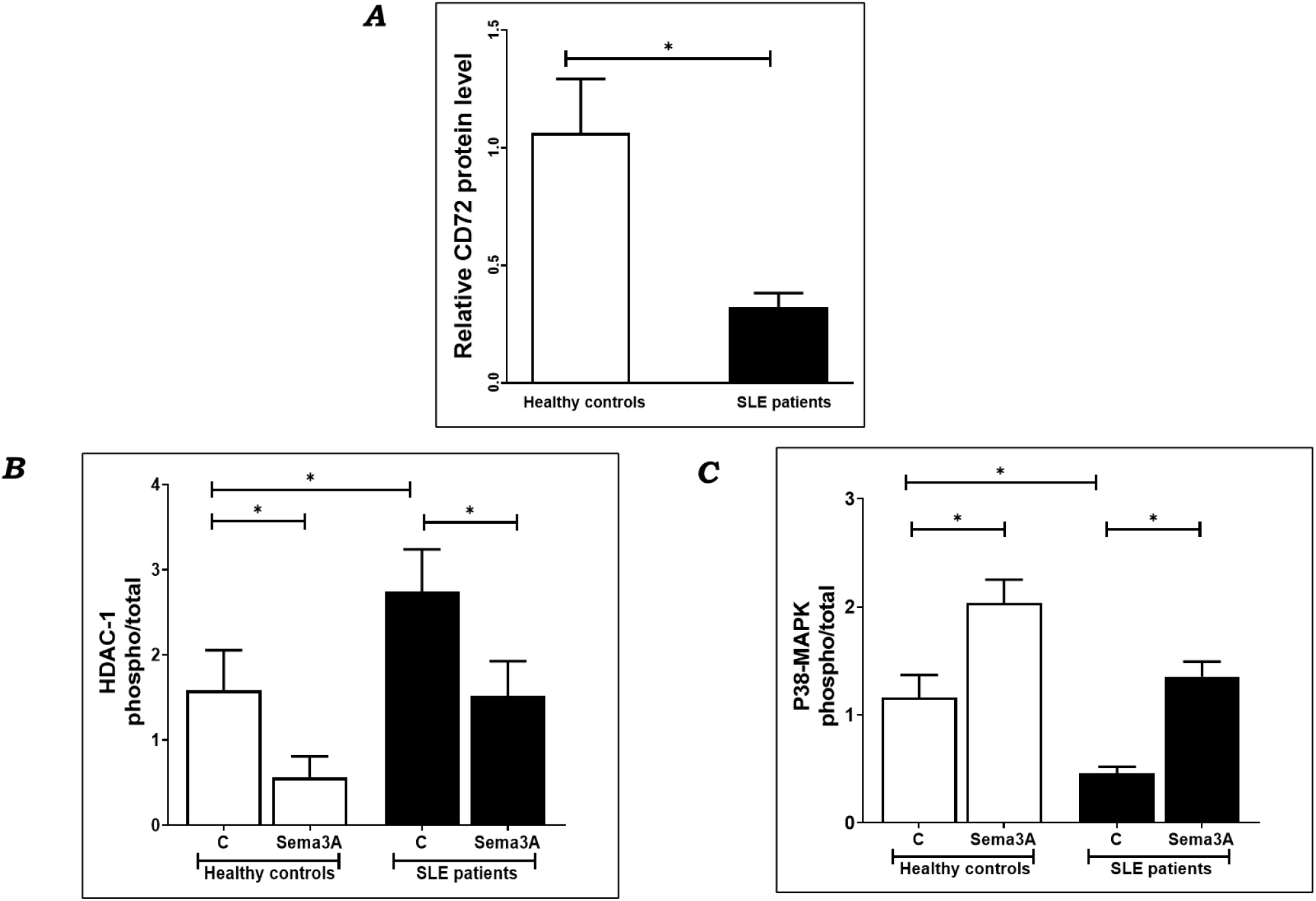
Sema3A binding to CD72 promotes the phosphorylation of p38-MAPK (Tyr182) and inhibits the phosphorylation of HDAC-1 (Ser421) in primary B cells from healthy controls and SLE patients. **(A)** Western blot showing of CD72 expression in activated B cells isolated from peripheral blood of healthy control and SLE patients. (CD72 expression levels were normalized to actin levels and quantified using ImageQuant TL Analysis software). **(B)** B cells purified from peripheral blood of SLE patients and healthy controls were activated with 1µM TLR9 agonist-CpG-ODN, and 5µg/ml CD40L at 37°C for 48 hours. After 48 hours, 10µg/ml of sema3A was added for 5 minutes at 37°C. Cell lysates were prepared in the presence of phosphorylation buffer as described. Blots prepared from the lysates were then probed with antibodies directed against phospho-HDAC-1 (Ser421) and phospho-p38-MAPK (Tyr182) and **(C)** as indicated. Shown are the resulting western blots and the quantification graphs which represent the mean± SD of the phospho/total ratio of each indicated component from 7 independent experiments.

### Generation of a modified truncated sema3A specifically binds CD72

Sema3A binding to the NRP-1 receptor initiates many complex responses such as changes in cytoskeletal organization and apoptosis of target cells *(20*-*23)*. Our observations suggest that a major part of the immune-suppressive response to sema3A is mediated through the activation of CD72 signal transduction and does not require NRP-1. We reasoned that if correct, then a modified sema3A able to activate CD72 but lacking the ability to activate NRP-1 signal transduction would have potential benefit for the treatment of autoimmune diseases. We have therefore produced a truncated sema3A that lacks the C-terminal NRP-1 binding domain downstream of amino acid 516 *(24, 25)*. In addition, because the active form of sema3A is a homo-dimer, and because the truncated C-terminal domain contains cysteines required for sema3A dimerization *(26)*, we exchanged a serine residue at position 257 with a cysteine residue to create an artificial dimerization site. Lastly, we added V5 and 6xHis epitope tags in frame upstream to a stop codon to facilitate the purification of T-sema3A (Fig. 7A). The cDNA encoding the truncated modified sema3A, which we named T-sema3A, was then expressed in HEK293 cells and purified using a nickel affinity column. We then verified that the purified T-sema3A is indeed able to form dimers since the active form of sema3A is a homo-dimer (Fig. 7B) *(23)*. To verify that T-sema3A is unable to activate NRP-1 mediated signal transduction, we performed cytoskeleton collapse assays. Human umbilical vein derived endothelial cells (HUVEC) respond to sema3A by contraction mediated by the NRP-1 receptor *(21, 23)*. Wild type sema3A was indeed able to induce the collapse of the cytoskeleton of the cells resulting in cell contraction. However, T-sema3A failed to induce cell contraction, demonstrating that T-sema3A is unable to activate NRP-1 mediated signal transduction (Fig. 7C). Stimulation of wild type BLCL cells with either sema3A or T-sema3A produced a relatively small induction of IL10 secretion (Fig. 7D). This small induction was likely the result of CD72 expression induced by sema3A and T-sema3A in these cells (Fig. 7E). Stronger stimulation of IL10 expression in response to sema3A or T-sema3A was observed in BLCL-CD72^+^ cells that express recombinant CD72 in addition to endogenous CD72 expression induced by sema3A or T-sema3A indicating that T-sema3A is able to induce the expression of an immune-suppressive factor such as IL10 in a CD72 dependent manner as well as wild type sema3A even though T-sema3A is unable to activate NRP-1 mediated signal transduction (Fig. 7D).

**Figure 7:**
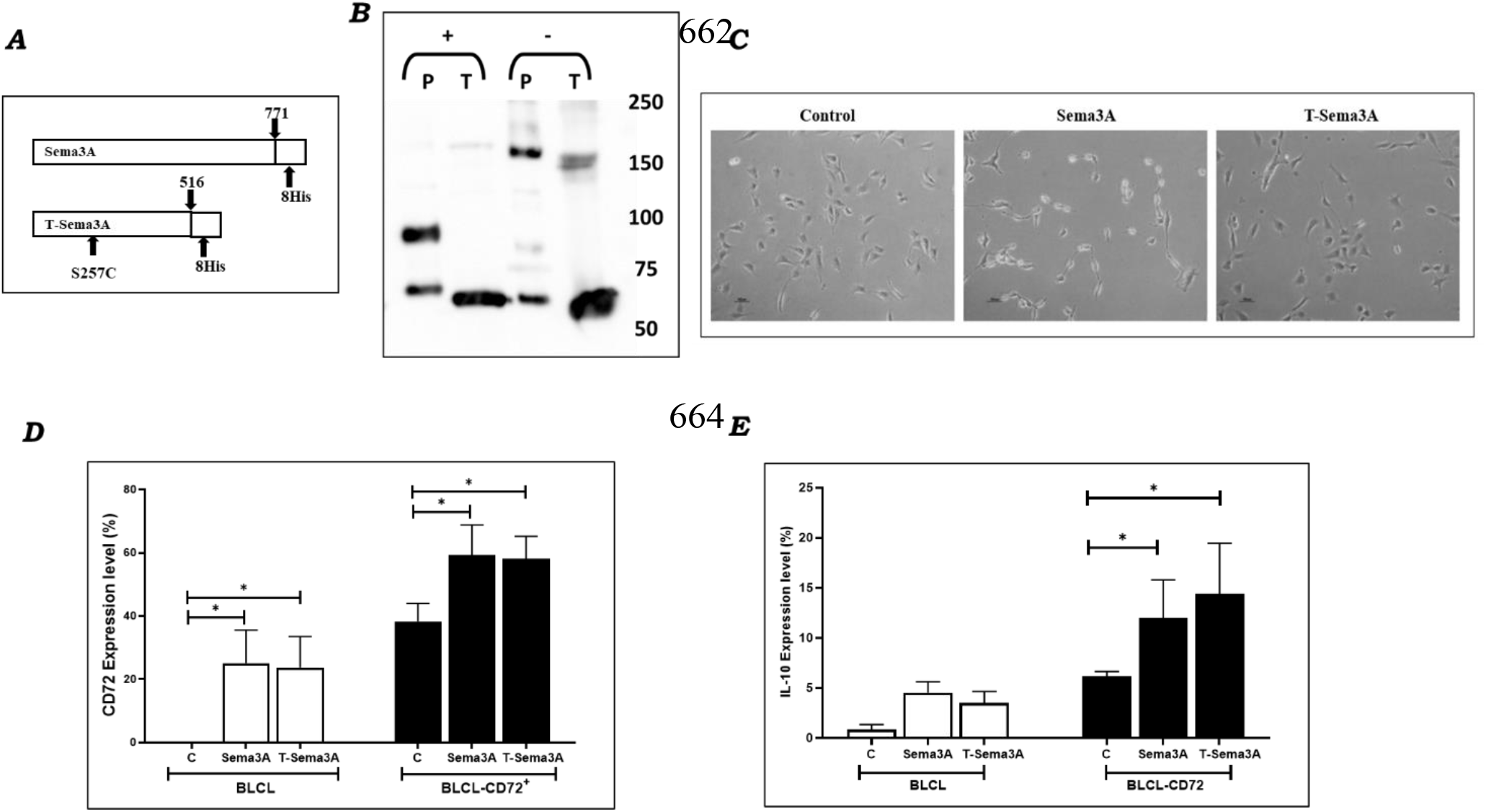
T(truncated)-sema3A. **(A)** The cDNA sequence of T-sema3A, a 1575bp construct truncated at base 1548 of the original sema3A with a point mutation at the 770 position, which exchanged a serine into a cysteine residue (s257c mutation), with 8xHIS epitope tags at the C terminal. **(B)** Ni-beads purified sema3A proteins (P=parental sema3A and T=truncated sema3A) were loaded in the presence of sample reducing agent (+) or without (-), and analyzed by western blot using an antibody directed against the N-terminal of sema3A. **(C)** Cytoskeleton collapse assay was performed on HUVEC cells in the presence of 200ng/ml of purified sema3A or T-sema3a or appropriate amount of elution buffer for 30 minutes at 37°C. After the incubation, the cells were photographed using a phase-contrast inverted microscope. (i) Round-shape cells is the natural morphology of the cells, (ii) as a result of sema3A, the cytoskeleton were collapsed, and the cells lost their round shape. (iii) T-sema3A has no effect on the round-shape cells. **(D)** BLCL and BLCL-CD72^+^ cells were stimulated with 10µg/ml sema3A or 10µg/ml T-sema3A or either carrier control for 24 hours at 37°C. CD72 and IL-10 **(E)** expression levels were evaluated using flow cytometry as described. (N=3).

## DISCUSSION

Sema3A is one of the best studied Semaphorins. It functions as a guidance factor for neurons as well as sprouting angiogenic blood vessels, and is one of the few Semaphorins that were characterized as immune semaphorins *(27)*. It is expressed by activated T cells, dendritic cells and endothelial cells, and is involved in the control of immune responses at all stages *(28)*. Following the binding of sema3A to its known NRP-1 receptor, NRP-1 associates with the plexin-A1 and the plexin-A4 receptors which do not bind sema3A independently, to form a functional sema3A signaling receptor in which all the three components are required *(29, 30)*. Activation of NRP-1 by sema3A induces the localized collapse of the cytoskeleton of target cells *(19)*. In immune cells sema3A was reported to down-modulate the activation of T cells and to enhance the expression of IL-10 in CD4^+^NRP-1^+^ T cells in NRP-1 dependent fashion *(31)*. In a later study we have found that stimulation of activated B cells (both of normal individuals and SLE patients) with sema3A is highly effective in the restoration of Bregs function and that this effect is associated with a sema3A induced increase in CD72 expression on their surfaces *(14)*. Sema3A also decreased the expression of TLR-9 in activate B cells of SLE patients suggesting that sema3A is able to reduce the auto-reactivity of B cells *(32)*. In agreement with these observations, we found that the survival of NZB/W mice was prolonged and the development of lupus nephritis was delayed following the injection of recombinant sema3A *(33)*.

Here we have found that sema3A function as a CD72 ligand and that the binding of sema3A to CD72 does not require the presence of NRP-1 since sema3A is also able to bind to CD72 in cells in which the NRP-1 gene was knocked-out. We also found that the stimulation of cells that express CD72 but no NRP-1 induces the phosphorylation of several intracellular signal transducing proteins such as PKC-theta and p38-MAPK and inhibits the phosphorylation of other secondary messengers such as STAT-4 and HDAC-1. These observations suggest that the binding of sema3A to CD72 induces sema3A dependent signal transduction separate from signal transduction mediated by the NRP-1 receptor. We have also observed that the inhibitory, CD72 mediated effects of sema3A are associated with a sema3A induced increase in the concentration of CD72 in B-cells. It is possible that the binding of sema3A inhibits the internalization and degradation of CD72 causing a net increase in its concentration and at the same time enabling efficient trunsduction of immune suppressive signals induced by sema3A. This will need to be examined in the future. These effects of sema3A are likely further enhanced by the inhibition of the binding of CD72 ligands that function as enhancers of immune responses such as sema4D/CD100 because both semaphorins seem to use a shared binding site on CD72. Thus, our findings suggest that the immune responses of B-cells are regulated by a sema4D/CD100-sema3A balance that is likely perturbed in auto-immune diseases such as SLE.

The secondary messengers whose phosphorylation state is modulated by sema3A via the CD72 receptor were all found to be of crucial importance in autoimmune diseases such as SLE. For example, IL-12 induced activation of STAT-4 in B cells contributes to the generation of autoantibodies and the secretion of inflammatory cytokines that play a role in many immune-mediated inflammatory diseases *(34)*. In agreement, STAT-4 deficient mice contain increased concentrations of CD4^+^Foxp3^+^ Tregs in their lymph nodes, suggesting that inhibition of STAT-4 expression contributes to Foxp3^+^ Tregs responses *(35)*. Histone deacetylases (HDAC) are transcription factors the demethylation and acetylation of which contributes to chromatin modifications in peripheral blood mononuclear cells. SLE patients were found to express higher levels of HDAC-1 transcripts as compared with healthy controls. This higher expression level was positively correlated with SLE disease activity and with the increased production of relevant auto-antibodies in SLE *(36)*. Our results suggest that the immune inhibitory effects of Sema3A may be mediated, at least in part, by CD72 mediated signal transduction that results in the inhibition of the phosphorylation of STAT-4 and HDAC-1 in B cells. Taken together these observations suggest that sema3A-CD72 signaling regulates crucial pathways in B cells that are important for the etiology of autoimmune and inflammatory diseases, and that modulation of this pathway by sema3A or T-sema3A may have potential therapeutic value for diseases such as SLE. We have also found that sema3A signaling mediated by CD72 induces the phosphorylation of PKC-theta and p38-MAPK. Both proteins were previously found to be involved in immune-regulatory mechanisms. P38-MAPK was observed to mediate IL10 expression induced by CpG in B cells and IFN-γ was found to induce the expression of the inhibitory cytokine IL-10 in B cells via the p38 and JNK signaling pathways *(37)*. Likewise, increased phosphorylation of PKC-theta is associates with increased secretion of the inhibitory cytokine IL-10 by T regs *(38)*. Thus, the binding of sema3A to CD72 in B cells seems to mediate, at least part of the immune inhibitory effects of sema3A in B-cells.

To summarize, our observations suggest that at least part of the immune inhibitory effects of sema3A on the activity of B-cells are mediated by the binding of sema3A to CD72. The binding seems to activate signal transduction cascades that lead to the phosphorylation and de-phosphorylation of secondary messengers that are known to play a role in the activation of B-cells. Our findings suggest that these activities do not require the presence of NRP-1, and further suggest that T-sema3A may perhaps be further developed as a therapeutic agent that selectively inhibits the activity of B-cells in autoimmune diseases such as SLE. These possibilities should be further examined in the future.

## MATERIALS AND METHODS

### Cells

- U87MG glioblastoma cells were obtained from the ATCC. U87MG-ΔNRP-1 cells in which the NRP-1 gene was knocked out were previously described *(19)*. U87MG-ΔNRP-1-CD72^+^ cells were generated from U87MG-ΔNRP-1 following infection with lentiviruses generated using a pLenti6.3/V5-DEST lentiviral vector directing the expression of full-length CD72 cDNA (Human CD72 9432bp sequence, clone 5226648, Dharmacon™) fused in-frame upstream of the stop codon with a V5 epitope tag (Gateway, Thermo Fisher Scientific).
- BLCL cell are Epstein-Barr virus-transformed primary B-lymphoblastoid cells (donor#213) and were obtained from Astarte Biologics, Inc. BLCL-CD72^+^ cells were generated from BLCL cells by infection with lentiviruses containing a pBABE-EGFP lentiviral expression vector (Gateway, Thermo Fisher Scientific) directing expression of the full-length CD72 cDNA.
- HUVECs (human umbilical vein derived endothelial cells) – primary endothelial cells isolated from the umbilical cord. These cells were isolated from fresh umbilical cords as previously described *(39)*. (These cells were not used beyond passage 8). Mycoplasma detection test was routinely performed in all cell lines using EZ-PCR™ Mycoplasma detection kit according to the manufacturer’s instructions (Biological Industries, Ltd).

### Real-Time PCR

QRT-PCR was performed using the StepOne Plus Real-Time PCR System with TaqMan Universal PCR Master Mix (2×), according to the manufacturer’s instructions (Applied Biosystems). The following TaqMan commercial primers were used: CD72 Hs00960066, NRP-1 Hs00826128, and RPLPO (Large Ribosomal Protein) Hs99999902 as an endogenous control gene. Data were analyzed by the Step One Plus Software using the Quantitation-Comparative method.

### Purification of sema3A

Conditioned medium of HEK293 cells producing recombinant sema3A fused with 6xHistidine tag in C-terminal was collected and purified using Ni-NTA agarose beads (QIAGEN). The protein activity was assessed using the cytoskeleton collapse assay *(19) (40)*.

### Cytoskeleton collapse assay

HUVEC cells were plated on gelatin-coated plates over-night at 37°C. At the day of the experiment, the cells were incubated with 200ng/ml of purified sema3A or T-sema3A or appropriate amount of elution buffer for 30 minutes at 37°C. Next, cells were photographed using a phase-contrast inverted microscope.

### Co-immunoprecipitation of CD72 and sema3A

U87MG as well as U87MG-ΔNRP-1 and U87MG-ΔNRP-1-CD72^+^ cells were incubated with sema3A for 20 minutes at 4°C followed by 5 minutes incubation at 37°C, then washed with ice-cold PBS, scraped, and lysed in the presence of lysis buffer (20mM Tris-HCl pH-7.5, 0.3M NaCl, 4mM EDTA, 1% Triton X-100). 1.5mg of total protein was incubated with anti-V5 tag mAb magnetic beads (MBL) for 2h at 4°C (or with anti-HA magnetic beads (Pierce™) as a control). The immunoprecipitates were subjected to SDS-PAGE and immunoblotted with anti-sema3A and anti-V5 antibodies. Bound antibodies were visualized using the EZ-ECL method (Biological Industries, Ltd), next the blots were viewed by Image Reader LAS4000.

### Purification of sema3A-Alkaline Phosphatase (AP)

Sema3A fused in frame with alkaline phosphatase (sema3A-AP) was purified from conditioned medium of HEK293-sema3A-AP cells (a gift from Prof. Oded Behar, *(41)*). The conditioned medium was concentrated using 30KDa Amicon Ultra centrifugal filter devices for a 50-fold concentration. The concentration was determined after InstantBlue Coomassie staining, with BSA as a concentration standard.

### Sema3A-alkaline phosphatase binding assay

U87MG as well as to U87MG-ΔNRP-1 and U87MG-ΔNRP-1-CD72^+^ cells were incubated with sema3A-AP for 1.5h at 4°C, followed by PBS wash and 20 minutes fixation with 4% paraformaldehyde, and 1h incubation at 65°C. Next, 5-bromo-4-chloro-3-indolyl phosphate and nitro blue tetrazolium (BCIP /NBT) liquid substrates for AP-enzyme (SIGMA) were added for overnight incubation at 4°C. The following day, a visible yellow-brown precipitate was microscopically demonstrated in case of AP-sema3A binding using a phase-contrast inverted microscope, and the mean color intensity per field (lum/µm) was analyzed using Image-Pro program.

For kinetics experiments, cells were incubated with an increasing concentration of unlabeled purified sema3A (0-100µg/ml) or recombinant human sema4D protein (0-50µg/ml) (Abcam) as competitive inhibitors, with a constant concentration of sema3A-AP (5µg/ml). The experiments were performed as was previously described. After using Image-Pro program for analysis, Graph Pad Prism software was used to draw the kinetics graphs and calculate the binding parameters (K_D_, Bmax, IC50).

### Co-immunoprecipitation of CD72 and SHP-1/2

BLCL-CD72^+^ cells were incubated with either sema3A or sema4D for 5 minutes at 4°C, then lysed in the presence of lysis buffer (50mM Tris-HCl pH-7.5, 150mM NaCl, 2mM EDTA, 2mM EGTA, 5mM NaF, 2mM Na_3_VO_4_, 10mM Na_4_P_2_O_7_, 1% Triton X-100). 1.5mg of total protein was incubated with protein G SureBeads™ (Bio-Rad) pre-coated with anti-CD72 monoclonal antibody (sc-25265) for 2h at 4°C. The immunoprecipitates were subjected to SDS-PAGE and immunoblotted with anti-SHP-1, anti SHP-2, anti-p-Tyr and anti-CD72 Antibody (sc-1707). Bound antibodies were visualized using the EZ-ECL method (Biological Industries, Ltd), next the blots were viewed by Image Reader LAS4000.

### Phosphorylation assay in BLCL and BLCL-CD72^+^ cells

BLCL-CD72^+^ cells were activated with 10µg/ml of sema3A for 5 minutes at 37°C. The phosphor array experiment was performed by following the antibody array user’s guide (Phospho Explorer Antibody Array (Full Moon-Catalog No: PEX100)), and the slides were shipped to the array-scanning unit of Full Moon Biosystems, Inc for scanning and image analysis.

For the results validation experiments BLCL and BLCL-CD72^+^ cells were lysis with phosphorylation lysis buffer (50mM Tris-HCl pH-7.5, 150mM NaCl, 2mM EDTA, 2mM EGTA, 5mM NaF, 2mM Na_3_VO_4_, 10mM Na_4_P_2_O_7_, 1% Triton X-100). 80µg of total protein were subjected to SDS-PAGE and immunoblotted with an antibody directed against a phosphorylated target protein, the blot was then stripped and re-probed with an antibody directed against total protein. Bound antibodies were visualized using the EZ-ECL method (Biological Industries, Ltd), next the blots were viewed by Image Reader LAS4000 and quantification of band intensity was performed using ImageQuant TL Analysis software.

### Phosphorylation assay in human primary B cells

#### Study population

SLE patients were evaluated by two rheumatologists (at the rheumatology unit, Bnai-Zion Medical Center, Haifa) for disease activity by the Systemic Lupus Erythematosus Diseases Activity Index (SLEDAI). The blood samples were taken before any therapy was initiated (steroid or cytotoxic drug). Informed consent was obtained from all studied individuals, and the study was approved by the local Helsinki committee for clinical studies.

Primary B cells were positively isolated from peripheral blood of healthy controls and SLE patients using anti-human CD22 micro beads, according to the manufacturer’s instructions (Milteny Biotec #130-046-401). The positively isolated B cells were activated with 1µM TLR9 agonist-CpG-ODN, and 5µg/ml CD40L at 37°C for 48 hours. After 48 hours, 10µg/ml of sema3A was added for 5 minutes at 37°C and the phosphorylation assay was performed as was previously described.

### Generation of Truncated-sema3A construct

To produce T-sema3A, the cDNA encoding the human sema3A was truncated at base 1751, A sequence encoding an epitope tag of 8 histidine residues was added upstream of a stop codon to enable purification on nickel beads. Also, a point mutation was introduced at base 770 counted from the ATG of T-sema3A which caused a serine residue replacement with a cysteine residue (S257C) in order to enable the formation of a T-sema3A cysteine bridge linked dimer.

### CD72 and IL-10 expression measurement in BLCL cells by flow cytometry

BLCL and BLCL-CD72^+^ cultured cells were treated with brefeldin A (420601, BioLegend) for 4hours, then seeded and stained with FITC Anti-Human CD72 antibody for 30 minutes. Next, cells were fixed, and permeabilized using FIX & PERM® cell Kit (GAS004, Invitrogen) and stained with APC anti-human IL-10 antibody for 30 minutes. The levels of IL-10 were evaluated in NAVIOUS EX flow cytometer (Beckman Coulter), and the results were analyzed using Kaluza, Analysis Software 2.1 (Beckman Coulter).

### Antibodies

Western blot antibodies:

- CD72 Antibody (G-5), sc-25265, Santa Cruz Biotechnology.
- CD72 Antibody (H-96), sc-1707, Santa Cruz Biotechnology.
- Neuropilin-1 Antibody (A-12), sc-5307, Santa Cruz Biotechnology.
- Sema3A Antibody, AF1250, R&D Systems.
- SHP-1 Antibody (C14H6), 3759, Cell Signaling Technology.
- SHP-2 Antibody, 3752, Cell Signaling Technology.
- p-Tyr Antibody (PY20), sc-508, Santa Cruz Biotechnology.
- HDAC1 Antibody, PA1-860, Invitrogen.
- p-HDAC1 Antibody (Ser421, Ser423), PA5-36911, Invitrogen.
- PKC theta Antibody (E-7), sc-1680, Santa Cruz Biotechnology.
- p-PKC theta Antibody (pT538.19), sc-136017, Santa Cruz Biotechnology.
- p38 MAPK Antibody, 9212, Cell Signaling Technology.
- pp38 MAPK (Thr180/Tyr182) Antibody, 4511, Cell Signaling Technology.
- Stat4 Antibody (C-4,) sc-398228, Santa Cruz Biotechnology.
- p-Stat4 Antibody (pY693.38), sc-136194, Santa Cruz Biotechnology.
- V5 Antibody Antibody, P/N 46-0705, Invitrogen.

Flow cytometry antibodies:

- FITC Anti-Human CD72 (316204), BioLegend.
- APC Rat Anti-Human IL-10 (554707) BD Pharmingen™.

### Statistical methods

Means obtained from each group were compared using one-way ANOVA followed by Kruskal-Wallis multiple comparison post-test. The following designations were used in the figures: *: p<0.05, **: p<0.01, ***: p<0.001, ****: p<0.0001 and non-specific: ns.

The statistical tests and the graphs were performed using Graph Pad Prism software.

## Funding

This work was supported by a grant from the Israel Science Foundation (to GN)

## Author contributions

Conceptualization: VZ, NE, GN

Methodology: VZ, NE, GN, JY

Investigation: VZ, NE, GN

Visualization: VZ, NE, SDA, KO, GN

Supervision: VZ

Writing – original draft: VZ. NG, NE

Writing – review & editing: VZ. NG, NE, KO

## Competing interests

Authors declare that they have no competing interests.”

## Data and materials availability

“All data are available in the main text or the supplementary materials.”

## Notes

### Competing Interest Statement

The authors have declared no competing interest.

